# Robust Colonization of Nasal Microbiota Across the Respiratory Tract and the Gut Modulates Innate Immune Response in a Germ-Free Pig Model

**DOI:** 10.1101/2025.10.31.685730

**Authors:** Shaurav Bhattarai, Tirth Uprety, Linto Antony, Chitra Sreenivasan, Sudeep Ghimire, Milton Thomas, David Francis, Steven Lawson, Feng Li, Joy Scaria, Radhey S Kaushik

## Abstract

The use of antibiotics in swine production has contributed to improved health and growth rates but also led to significant challenges, including the emergence of antibiotic-resistant bacteria and disruptions in the microbiome. This study employs a germ-free pig model to investigate the transplantation of the nasal microbiome into the respiratory tract, exploring its potential to colonize multiple body sites and its impact on host immunity. Using germ-free piglets inoculated with nasal microbiota from healthy donors, we examined the dynamics of microbial community formation across the respiratory and gastrointestinal tracts and assessed the effects of antibiotic intervention. Our findings reveal that specific taxa, predominantly from the phyla Proteobacteria and Firmicutes, cross colonized both respiratory as well as gastrointestinal tract. This cross-colonization highlights the adaptability of nasal microbiota and suggests a significant role in modulating host immunity through the gut-lung axis. The administration of the antibiotic tulathromycin on day 11 post-inoculation altered both nasal and gut microbiota compositions, reducing microbial diversity and impacting immune gene expression and cytokine profiles. These results underscore the complex interactions within the gut-lung axis and emphasize the implications of antibiotic use on microbial health and host immunity. Understanding these interactions provides crucial insights into alternative strategies for managing health in swine production without relying on antibiotics. Our study advocates for a reevaluation of antibiotic practices and supports the development of microbiome-centered interventions in swine health management.

## Introduction

The nasal microbiome is a diverse community of microorganisms inhabiting the nasal cavity and plays an important role in maintaining the health and well-being of farm animals. The nasal microbiome serves as a barrier against pathogenic microbes, aids in developing innate and adaptive immune systems, and modulates inflammatory immune responses (1–5). A healthy nasal microbiome can prevent pathogen colonization, thereby reducing the risk of respiraory infections and perturbations in the nasal microbiome can increase susceptibility to respiratory infections leading to poor animal health and reduced production (3). The nasal microbiome modulates the immune responses by interacting with the immune cells and altering the expression of immune-related genes in the respiratory tract (6). Studies have shown that pigs with high immune competencies are enriched with beneficial bacteria such as Lactobacillus in the respiratory tract, which contributes to generating an effective immune response to respiratory pathogens. Similarly, enriched Lactobacillus and Prevotella in the nasal microbiome have been associated with increased production performance in pigs by modulating nutrient utilization, growth rate, and immune functions (1, 7).

The composition of the nasal microbiome is influenced by several factors, including age, environment, genetics, vaccination, and housing practices (8–10). Systemic antibiotics administration alters the microbiome’s composition at mucosal surfaces, including the gut and respiratory tract, but their impact on the expression of various innate immune genes is not well studied (11, 12). Identifying the nature of bacterial community shift during antibiotics treatment can help to develop effective strategies for promoting swine health and production (13, 14).

The gnotobiotic model is a valuable tool for studying the role of the microbiome in the health and disease and these gnotobiotic pigs are raised in germ-free environments and lack microbial colonization. This model allows precise control over the microbial environment which facilitates the study of the impact of individual or complex microbial communities on the host (15, 16). The gnotobiotic pig model has been used to evaluate the virulence and pathogenesis of several pathogenic microbes, including rotavirus (17), reovirus (18), influenza virus (19), Clostridioides difficile (20), porcine epidemic diarrhea virus (21) and porcine reproductive and respiratory syndrome virus (22, 23).

While the gut microbiome has received more attention and has been extensively studied, only a few studies have focused on the nasal microbiome composition and its effects on immunity (4, 24) . We hypothesize that inoculation of the nasal microbiome in germ-free pigs upregulates innate immune responses in the respiratory tract and establishes a stable microbial community in the respiratory tract and gut. In this study, we developed a gnotobiotic pig model to study bacterial colonization after nasal microbiome inoculation and evaluated the shift in microbial community after antibiotics treatment. Furthermore, we also studied innate immune gene expression at the nasal mucosa and surrounding lymphoid organs. We found that nasal microbiome inoculation upregulates the expression of various Toll-like receptors, chemokines, and inflammatory cytokines leading to increases in lymphocyte infiltration in the nasal mucosa. Similarly, we characterized a stable nasal and gut microbiome following nasal microbiome inoculation and observed a shift in gut and nasal microbiome after Tulathromycin antibiotics treatment.

## Materials and Methods

### Ethical Statement

All the animal experimental procedures were approved by the Institutional animal care and use committee (IACUC) of South Dakota State University (SDSU).

### Nasal inoculum preparation

Nasal swabs were collected from two to three-week-old conventional piglets reared in the SDSU swine unit. We collected two nasal swabs (one from each nostrils) per animal and a total of 50 animals were used. Swabs were collected after cleaning the outside of the nostrils and transferred immediately to a tube containing Brain Heart Infusion (BHI) broth. After vigorous vortexing, all samples were pooled together to prepare homogenous nasal microbiome inoculum. Bacterial pellets were obtained after centrifuging the inoculum at 7000 rpm for 10 minutes. Following resuspension of the pellets in BHI broth, 10% DMSO stocks were prepared, and inoculumwase stored at -80^0^C until further use. Prior to the inoculation in gnotobiotic pigs, aliquot of frozen stock was thawed and serially plated in BHI plates both aerobically and anaerobically to assess the bacterial load in the sample.

### Animal experiment

Sow close to parturition was purchased, and a licensed university veterinarian performed C-section to obtain the germ-free piglets. An appropriate setup was built to avoid exposure of the newborn piglets to non-sterile conditions. 12 piglets were kept randomly in three separate pens with four piglets in each cell. Physical contact between the piglets was avoided, and animals were fed separately. Twenty-five ml of commercial bovine colostrum preparation was offered daily to the piglets for the first four days. Serum was extracted from the sow to provide passive immunity to the piglets. 15ml of the sow serum was injected intraperitoneally for seven days, starting from the second day. On the 4^th^ day, eight piglets from 2 pens received intranasal inoculation of 1.44 *10^5 CFU of the nasal microbiome. Similarly, four piglets from the remaining pen were mock-inoculated with sterile PBS. After 11 days of the inoculation (dpi), four piglets from one of the nasal microbiome inoculated groups received an intramuscular injection of tulathromycin antibiotics at the dose of 2.5mg/kg body weight. Piglets from all the groups were euthanized seven days after the antibiotic’s treatment (18dpi). Blood samples were collected before the nasal microbiome inoculation (0 dpi), 11 dpi, and 18dpi for the quantification of serum immunoglobulins. Nasal and fecal swabs were collected before the inoculation, 6dpi, 11dpi, and 18dpi. Nasal mucosa, palatine tonsil, mediastinal lymph node, lung, and PBMC were collected at necropsy.

### Microbial DNA preparation

Microbiome DNA was extracted from the frozen aliquot of pooled inoculum as well as the nasal and fecal swabs collected from individual animals at various time points. Frozen samples were thawed and were vigorously vortexed with 1 ml of PBS. The suspension was then centrifuged at 7,000 rpm for 10 minutes to pellet the bacteria. The resulting pellet was utilized for DNA extraction using the DNeasy PowerSoil Kit (Qiagen) according to the manufacturer’s protocol. Briefly, bacterial cells were lysed chemically using an SDS-based lysis solution and mechanically through bead beating. This step was followed by the removal of contaminating organic and inorganic materials, including cell debris and proteins. A highly concentrated salt solution was used to precipitate the total genomic DNA, which was then transferred to a silica membrane in a spin column. After a two-step, on-column cleaning process, the DNA was eluted with ultra-purified nuclease-free water. The quality of the DNA was assessed using NanoDrop (Thermo Fisher Scientific) and the DNA was stored at -20°C until further analysis.

### Targeted amplicon sequencing and community profiling

The microbiome community composition of all samples were determined by sequencing the V3- V4 region of the 16s rRNA gene using Forward Primer - 5’ CCTACGGGNGGCWGCAG 3’, and Reverse Primer - 5’ GACTACHVGGGTATCTAATCC 3’ (Illumina, Inc.). Following a bead-based cleaning of the amplicons using AMPure XP beads (Beckman Coulter), sequencing library was prepared using Nextera XT library preparation kit (Illumina, Inc.). We used a dual indexing method for multiplexing the samples. Sequencing of the pooled library was then performed on the Illumina Miseq platform using paired end V2 chemistry (250bpx2).

We used FastQC (https://github.com/s-andrews/FastQC ) to check the quality of the reads as well as for demultiplexing and adaptor-trimming. Samples that passed the quality check were analyzed using Qiime 2 microbiome analysis pipeline(25). Read quality filtering and feature table construction after removing chimeric sequences were performed using the DADA2 pipeline(26). After denoising, the taxonomic assignment was performed in Qiime2 using Greengene database(27). Statistical and visual exploration of feature abundance data generated in qiime2 was performed using MicrobiomeAnalyst(28).

### ELISA and H&E staining

Quantification of total serum IgA, IgG, and IgM was performed using a commercially available ELISA kit (E1101-102, E1101-104, and E1101-117) following the manufacturer’s instructions.

Formalin fixed paraffin embedded (FFPE) nasal mucosa tissue were sliced in 5 µm thickness using microtome and mounted in lysine coated glass slides. H&E staining was performed as explained elsewhere (29). Briefly, sections were deparaffinized using xylene and rehydrated using graded ethanol. Following rinsing with distilled water, sections were stained with hematoxylin and eosin solution. Stained sections were dehydrated in graded ethanol and observed under Olympus upright microscope. Nasal mucosal layer tissue morphology and cellular details were recorded and analyzed.

### RNA isolation, cDNA preparation, and qPCR

Total RNA was isolated from nasal mucosa, palatine tonsil, mediastinal lymph node, and lungs using the trizol method. Briefly, 100 mg of the snap-frozen tissue sample was used to isolate the RNA. Nanodrop 2000 was used to quantify the extracted RNA. For the reverse transcription, 1ug of the RNA was used to synthesize cDNA following the kit manufacturer’s instructions. (Applied Biosystem, N8080234). The Oligo dt method was applied to reverse transcribe mRNA from the total RNA selectively. Prepared 20ul cDNA was diluted 5x in nuclease-free water.

Primers specific to porcine TLRs (1,2,3,4,5,6,7,8 and 9), RIG-I, MDA-5, IL-8, MCP-1, IL-1a, IL-1b, IL-6, TNF-a, IL-10, INF-a, and INF-b was used to study the gene expression profile in the tissues (Table 1). 2ul of the diluted cDNA was used as a template. 1ul each of forward and reverse primers (10uM), 10ul of SYBR green master mix (Qiagen, 330523), and 6ul of nuclease- free water were used for the qPCR reaction. The ribosomal protein L 4 (RPL4) gene was used as an internal control. Delta CT was calculated and used for the statistical analysis. Thermal cycles were as follows: initial denaturation at 95^0^c for 10 minutes, 40 cycles of denaturation at 95^0^c for 15 seconds, annealing for 30 seconds, and final extension at 72^0^c for 30 seconds. A melt curve was obtained to confirm the specificity of the qPCR reaction. qPCR was performed using Quantstudio^TM^ 6 Flex Real-Time PCR system (Applied Biosystem, NJ, USA).

**Table 1:**
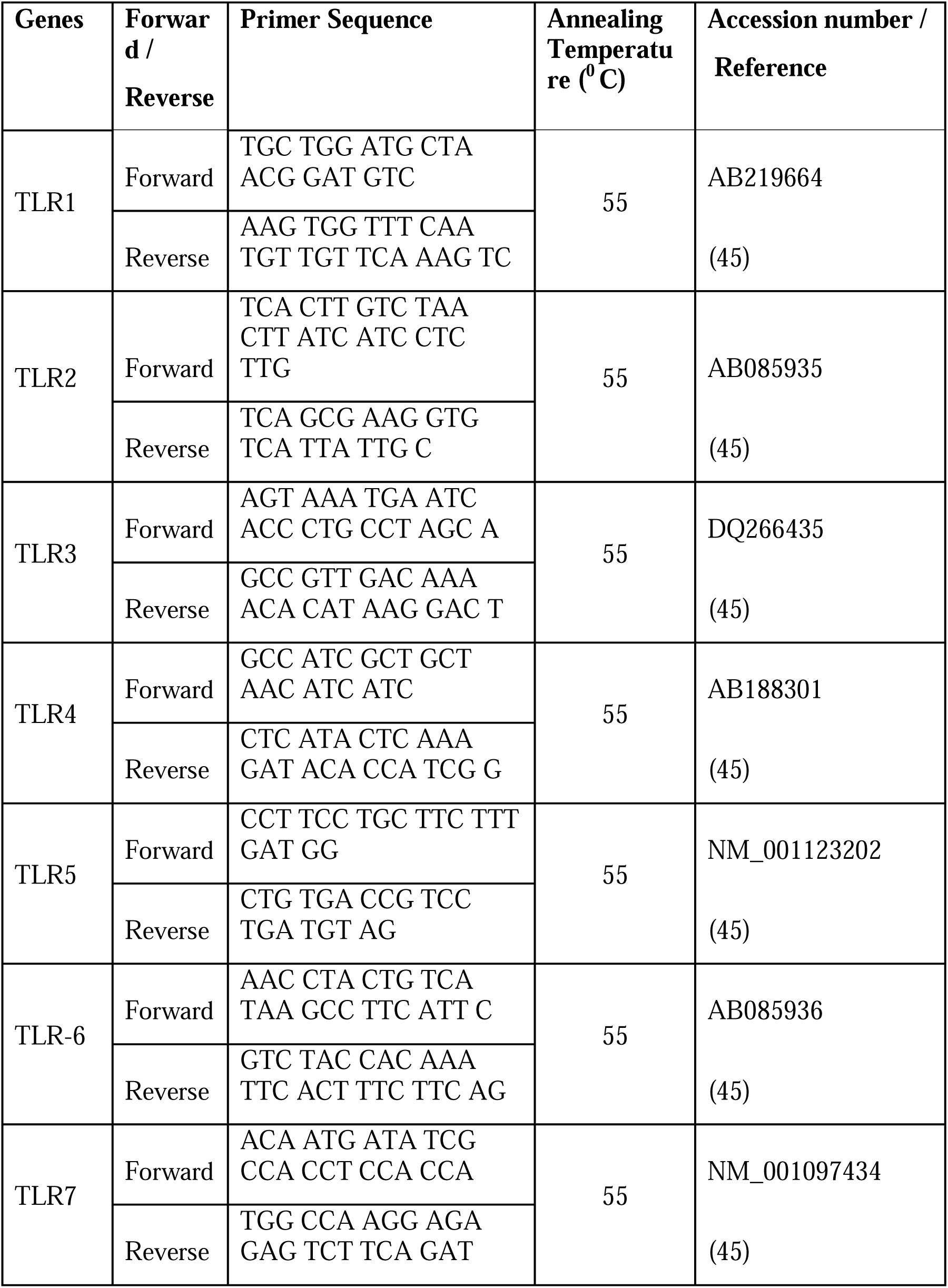
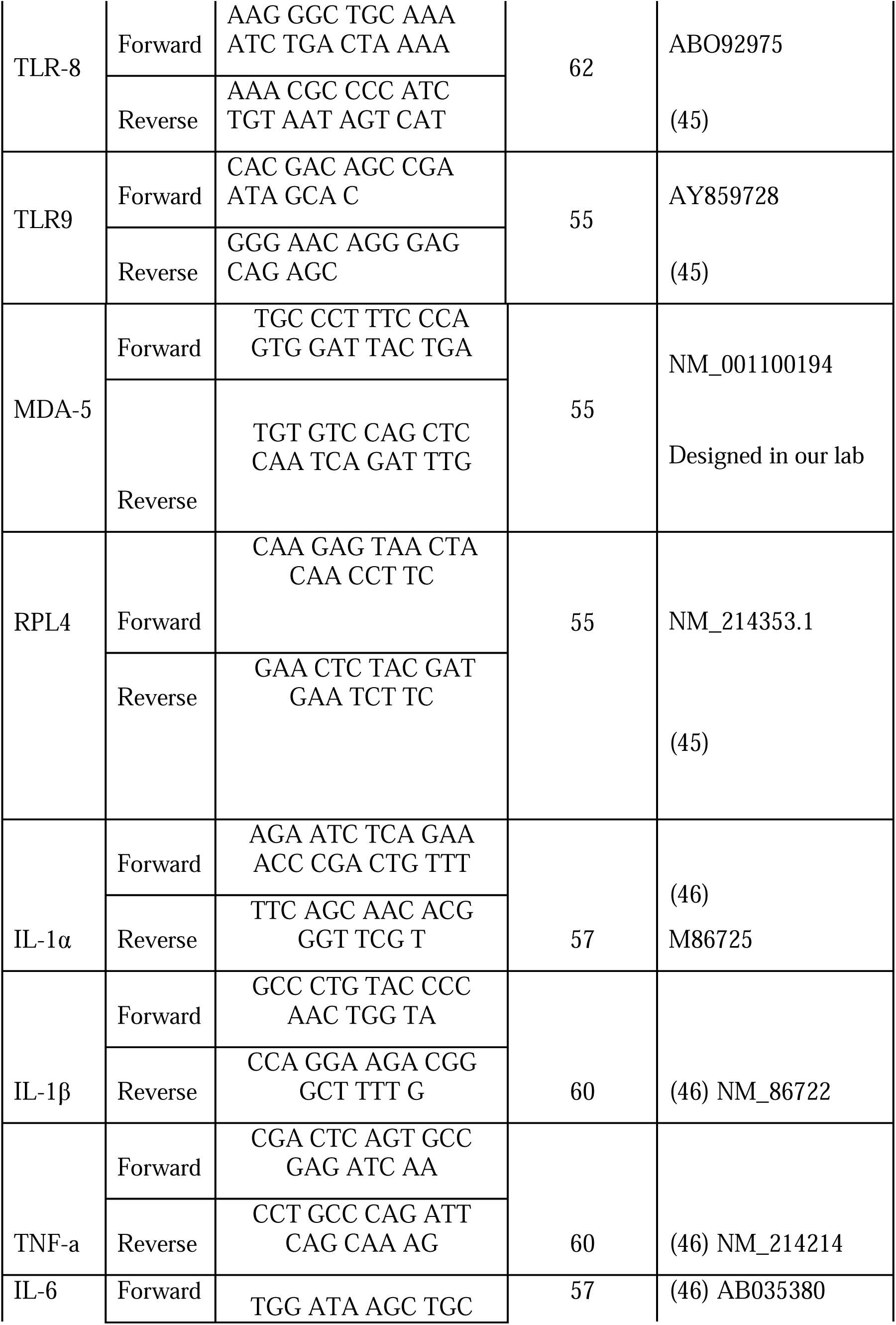
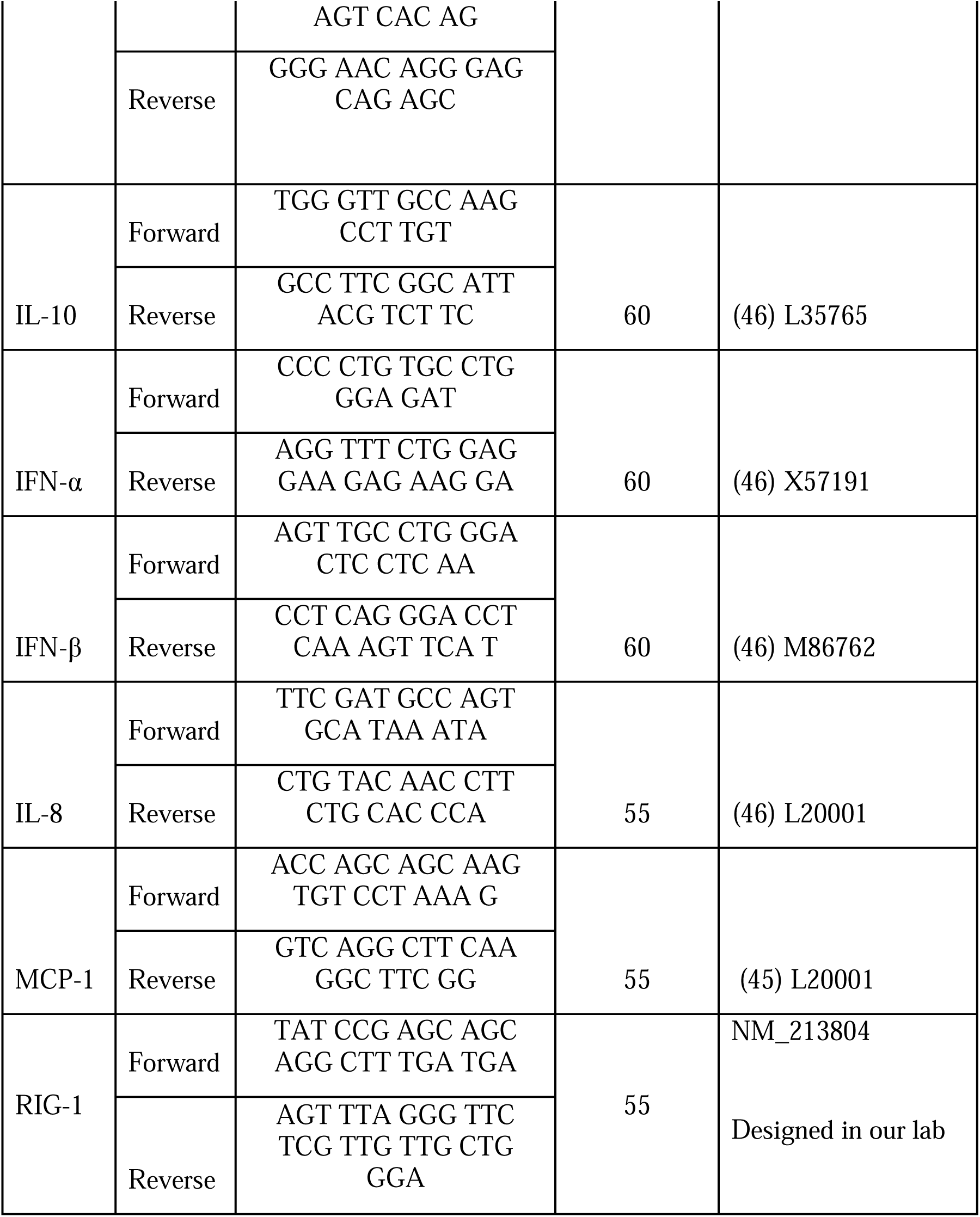
List of primers used in the study.

### PBMC proliferation assay

Fifty thousand live PBMC cells were seeded in RPMI media supplemented with 10%FBS and 1% antibiotics (penicillin-streptomycin). Cells were cultured in 96-well flat button plates and incubated at 37^0^c with 5% CO2. After 24 hours of seeding, cells were stimulated with either Con-A, LPS, or PHA at 10ug/ml concentration or left unstimulated. Cell proliferation assay was performed using a commercially available BrdU kit (Roche, 11647229001) following the manufacturer’s instructions. After 72 hours of stimulation, 10ul of BrdU labeling solution was added to all the wells at the final concentration of 10Um of BrdU per well and incubated overnight. Media was removed, and cells were dried at 60^0^c for 1 hour. Following fixation, cells were stained with an anti-BrdU antibody. The color was developed using the kit substrate, and the reaction was stopped by adding 25ul of 1M H_2_SO_4_. Absorbance was measured at 450nm wavelength in a synergy plate reader. Background absorbance was determined with unstimulated BrdU-stained cells.

### Statistical analysis

One-way ANOVA and Tukey’s post-hoc test was used using Rstudio (Version 1.4.1106) to evaluate the significance of qPCR data. The Shapiro-Wilk test was used to test the normality of the samples and all the values were found to be normally distributed. For serum antibody quantification and PBMC proliferation assay, a student-test test was applied using Microsoft excel. A p-value of less than 0.05 was considered statistically significant.

## Results

### Dynamics and impact of nasal microbiota transplantation on germ-free piglets: Colonization patterns across respiratory and gastrointestinal tracts

In this study, we utilized a germ-free piglet model to examine the dynamics of nasal microbiome colonization and its impact on host immunity. Since germ-free piglets are delivered by cesarean section and reared in sterile isolators, they do not receive maternal immunoglobulins, making them potentially more susceptible to the invasive effects of commensal microbes. Our pilot experiments confirmed this increased susceptibility, as evidenced by some mortality among the germ-free piglets inoculated with nasal microbiota from healthy piglets. To address this issue, we implemented a strategy to provide passive immunity to the piglets. This strategy comprised of two approaches: initially, piglets were administered sterile bovine colostrum orally for xx days. Subsequently, on days 3 and 6, piglets received sow serum intraperitoneally. This approach successfully protected the piglets, with no further mortality observed. After inoculating with nasal microbiota, we monitored the piglets for 18 days to assess the successful colonization and assembly of the microbial community, as well as immune modulation.

We assessed the colonization and trajectory of microbiome community formation in both nasal and fecal samples through bacterial 16S rRNA gene sequencing-based community profiling. The pooled nasal sample used for microbiota transplantation was also sequenced and compared against the gnotobiotic community to evaluate differences in bacterial community structure between donor and recipient microbiota. At the phylum level, Proteobacteria, Bacteroidetes, Firmicutes, and Actinobacteria dominated the donor nasal microbiome (Supplementary Figure 1). At the family level, Pasteurellaceae, Weeksellaceae, and Moraxellaceae accounted for about 80% of the abundance, with the remaining 20% comprising Streptococcaceae, Clostridiaceae, Micrococcaceae, Ruminococcaceae, Lactobacillaceae, and Lachnospiraceae (Figure 1A). We analyzed the nasal microbiome composition in piglets on days 6, 11, and 18 and compared it to that of the inoculum. The overall nasal microbiome community composition was largely preserved following implantation in ex-GF piglets, albeit with some changes. The primary difference was the expansion of the phylum Tenericutes and a reduction in Bacteroidetes. The primary contributor to this change was the expansion of Mycoplasmataceae, which was very low in abundance in the donor microbiota. This family showed a gradual increase in abundance over time and became the most abundant in the nasal microbiota by day 18 (Figure 1A). Conversely, Bacteroidetes, the second most abundant component of the donor nasal microbiota (Supplementary Figure 1), exhibited a reverse trend. The family Weeksellaceae, representing the phylum Bacteroidetes in the donor microbiota and accounting for 32.9% of the total abundance, was reduced to less than 1% by day 18 post-inoculation (Figure 1A). Despite these differences in relative abundances at various taxonomic levels, group-wise alpha diversity measurements using the Shannon index indicated that by day 18, the richness and evenness of the nasal microbiota community within a sample were becoming more similar to those of the donor microbiota (Supplementary Figure 2).

**Figure 1:**
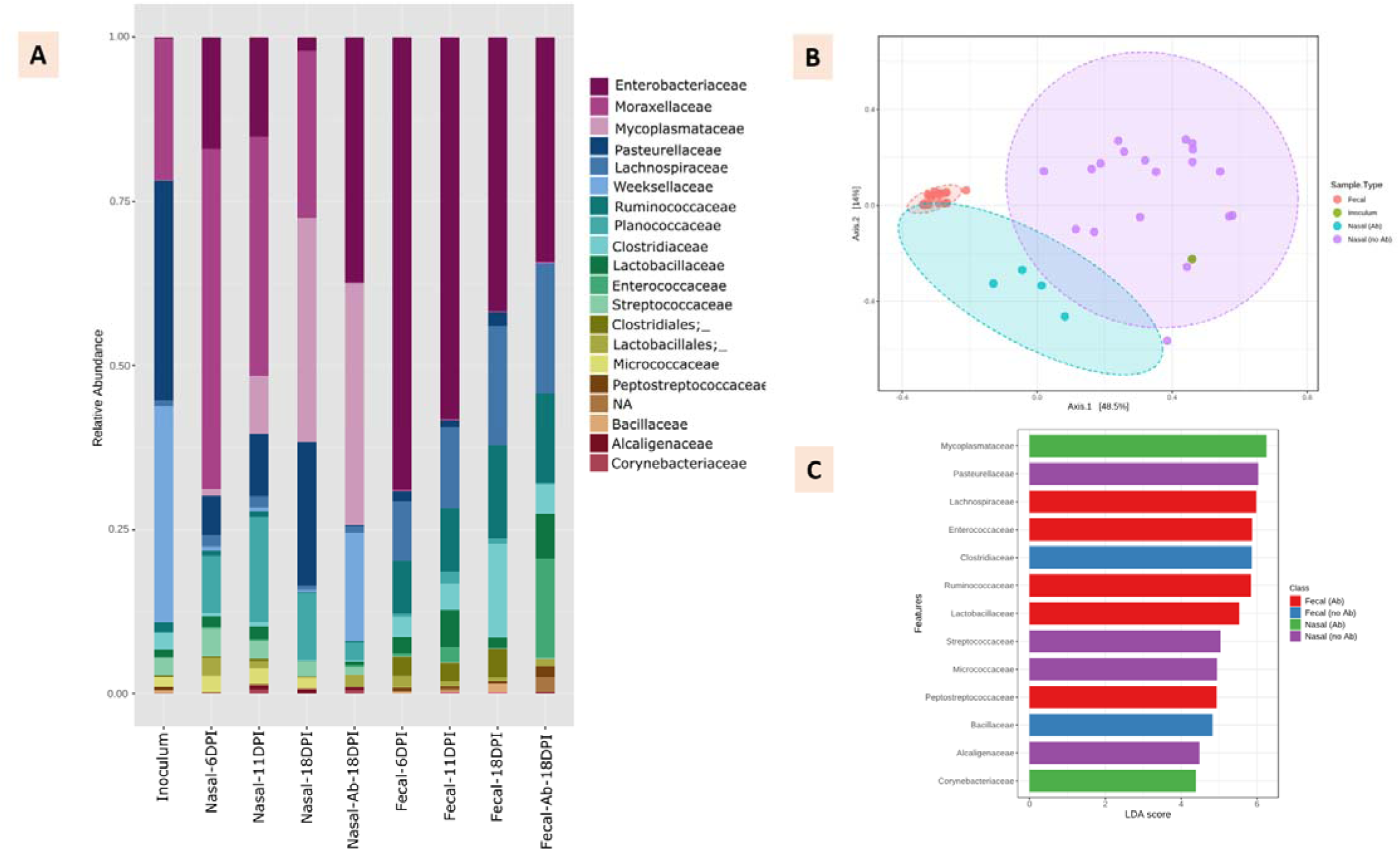
Transplantation of nasal microbiota from conventional piglets resulted in distinct bacterial community formation in the nasal cavity and gut of germ-free piglets. Bar plot (A) represents the abundance of top 20 bacterial families in both nasal and in the fecal samples collected at 6th, 11th, and 18th days after post inoculation (DPI) in comparison to pooled donor nasal microbiota (inoculum). PCoA plot (B) generated based on Bray-curtis index shows treatment with antibiotic (Ab) via nasal route resulted in the shift of nasal community. LEfSe analysis (C) at the family level shows certain bacterial features (Log LDA score > 4) are significantly present in nasal, with (Ab) and without antibiotic (no Ab), as well as in fecal community.

Next, we assessed whether, in addition to colonizing the upper respiratory tract, the nasal microbiota could also stably colonize the gastrointestinal tract of the germ-free piglets, potentially modulating host immunity. When fecal samples were analyzed, we indeed found this to be the case (Supplementary Figure 1). However, there were substantial differences between the nasal microbiome community and the gut microbiome community. While members of the phyla Proteobacteria and Firmicutes colonized both the respiratory tract and the gut, Bacteroidetes were completely absent in the gut. Members of the families Clostridiaceae and Bacillaceae were significantly associated (FDR-adjusted p-value < 0.05) with the fecal community by the 18th day post-inoculation (DPI), as determined by LEfSe analysis (Figure 1C).

### Antibiotic alters the nasal as well as gut microbiota community structure

Antibiotic exposure is one of the main factors that disrupt the microbiota structure and shift the bacterial community to an alternative stable state. To understand the perturbation elicited by antibiotics on the establishing nasal microbiota in the gnotobiotic piglets, we used the antibiotic tulathromycin, an efficient macrolide antimicrobial agent for swine respiratory diseases. We separated one group of gnotobiotic piglets on the 11^th^ day post introduction of nasal microbiota and treated them with a single dose of tulathromycin at a dose rate of 2.5 mg/kg body weight.

We observed significant reduction of alpha diversity as measured by the Shannon index compared to that of the non-antibiotic treated group as well as the donor nasal microbiota (Supplementary figure 2). Differences in the community composition between pre- and post-antibiotic nasal samples was evaluated using the Bray-Curtis dissimilarity index. Post-antibiotic samples showed a significant (PERMANOVA, p-value < 0.001) shift in the nasal microbiota community from the pre- and non-antibiotic treated counterparts (Figure 1B). Compared to the non-antibiotic treated group, any drastic elimination effect on the microbiota composition at the phylum level was not found instead increased the relative abundance of the family Weeksellaceae and thus the phylum Bacteroidetes (Figure 1A, Supplementary figure 1).

Antibiotic exposure caused an increase in the relative abundance of Enterobacteriaceae and Weeksellaceae, while a profound reduction occurred in Pasteurellaceae and Moraxellaceae (Figure 1A). However, LEfSe analysis on endpoint samples (18^th^ DPI) showed a significant (FDR adjusted p- value < 0.05) association of only Mycoplasmataceae and Corynebacteriaceae families with log LDA score of more than 4.0 (Figure 1C).

### Microbiome colonization modulates host immunity

To determine the effect of microbiota colonization on host immunity, we quantified total serum immunoglobulins and profiled cytokines, chemokines and TLRs. In the case of IgG, no significant differences were observed between the treatment groups throughout the experimental period; total serum IgG levels remained relatively constant across all three groups (Fig 2a).

**Figure 2:**
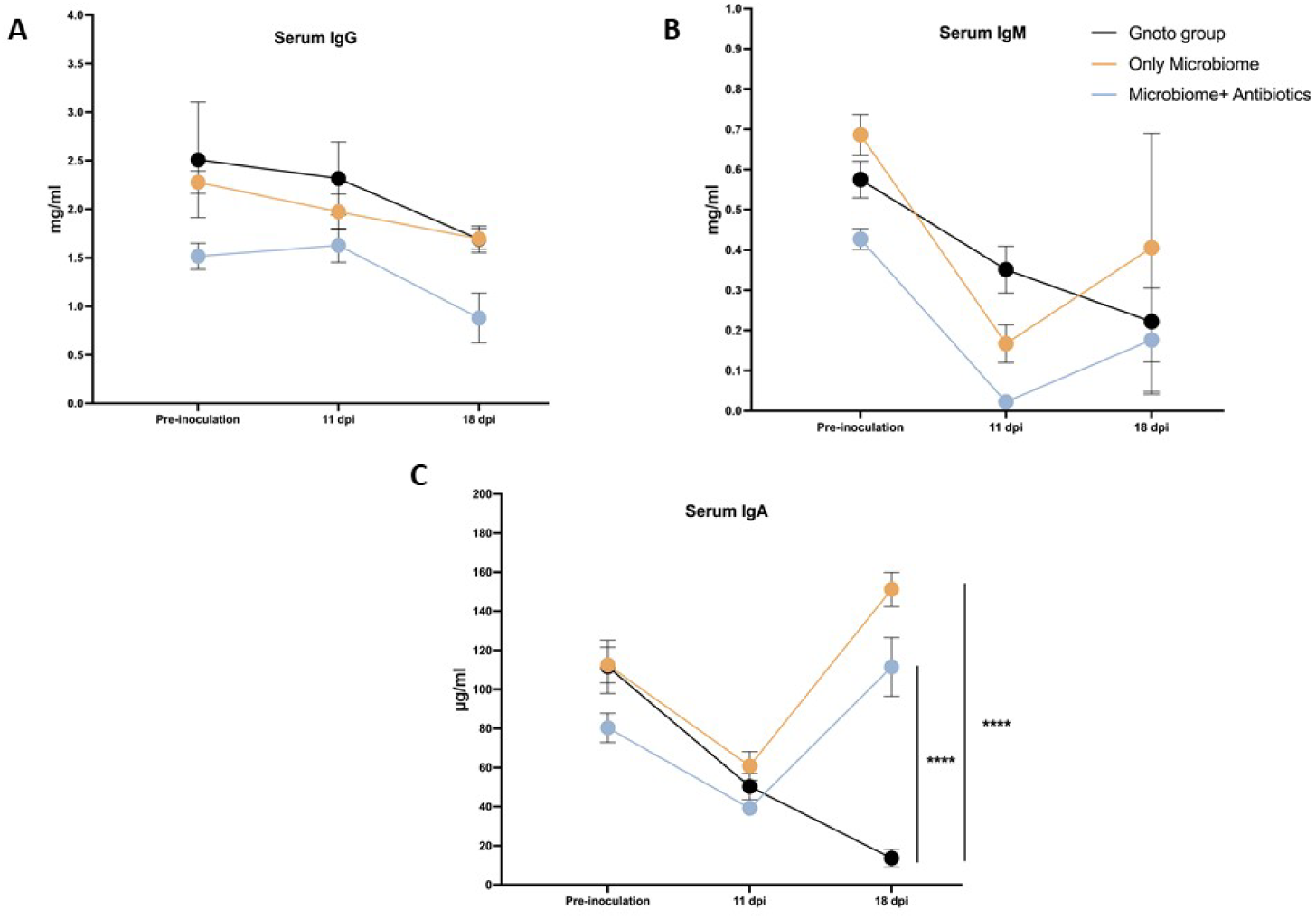
Total serum antibodies level in germ-free (gnotobiotic) group, microbiome plus antibiotics treated, and microbiome only inoculated groups. Figures 1a, 1b, and 1c show total serum IgG, IgM, and IgA expression, respectively. Three-time points on X-axis represent serum antibodies level of pre-nasal microbiome inoculation (0dpi), 11 days post-inoculation, and 18 days post-inoculation. Y-axis represents the units, IgG and IgM are plotted as mg/ml unit, and IgA is plotted using µg/ml unit. The error bar represents the standard error of the mean (SEM). Asterisk (*) denotes statistically significant, and a p-value of less than 0.05 was considered statistically significant.

Similarly, total serum IgM levels did not show significant changes between the treatment groups (Fig 2b). All groups exhibited a decrease in total serum IgM levels at 11 days post-inoculation. By 18 days post-inoculation, serum IgM levels increased in both the microbiome plus antibiotics and microbiome-only treated groups; however, the levels continued to decrease in the gnotobiotic group. Despite these fluctuations, the changes in IgM levels were statistically non-significant.

Interestingly, significant differences in serum IgA levels were observed at 18 days post-inoculation (dpi) between the germ-free group and the nasal microbiome inoculated groups (Fig 2c). At 11 dpi, total serum IgA levels were lower in all three groups (50.32 ± 6.26 µg/ml, 39.22 ± 3.65 µg/ml, and 60.84 ± 7.36 µg/ml, respectively) compared to the pre-inoculation levels (111.59 ± 13.6 µg/ml, 80.35 ± 6.48 µg/ml, and 112.49 ± 9.15 µg/ml, respectively). By 18 dpi, IgA levels decreased further in the germ-free group (13.66 ± 4.61 µg/ml). In contrast, IgA levels significantly increased in both the microbiome plus antibiotics treated group (111.51 ± 15.03 µg/ml) and the microbiome-only treated group (148 ± 11.68 µg/ml), indicating that the inoculated nasal microbiome induced IgA production in the piglets.

Immune modulation was further assessed by quantifying the expression levels of various Toll-like receptors (TLRs), chemokines, and cytokines in the nasal mucosa, palatine tonsil, mediastinal lymph node, and lung tissue. In the nasal mucosa, expression of selected TLRs was upregulated in groups inoculated with nasal microbiome and those treated with antibiotics compared to the germ-free control groups (Figure 3). Specifically, the expression of TLR2 and TLR4 was significantly higher at the cell surface in both microbiome-treated groups (Fig 3a).

**Figure 3:**
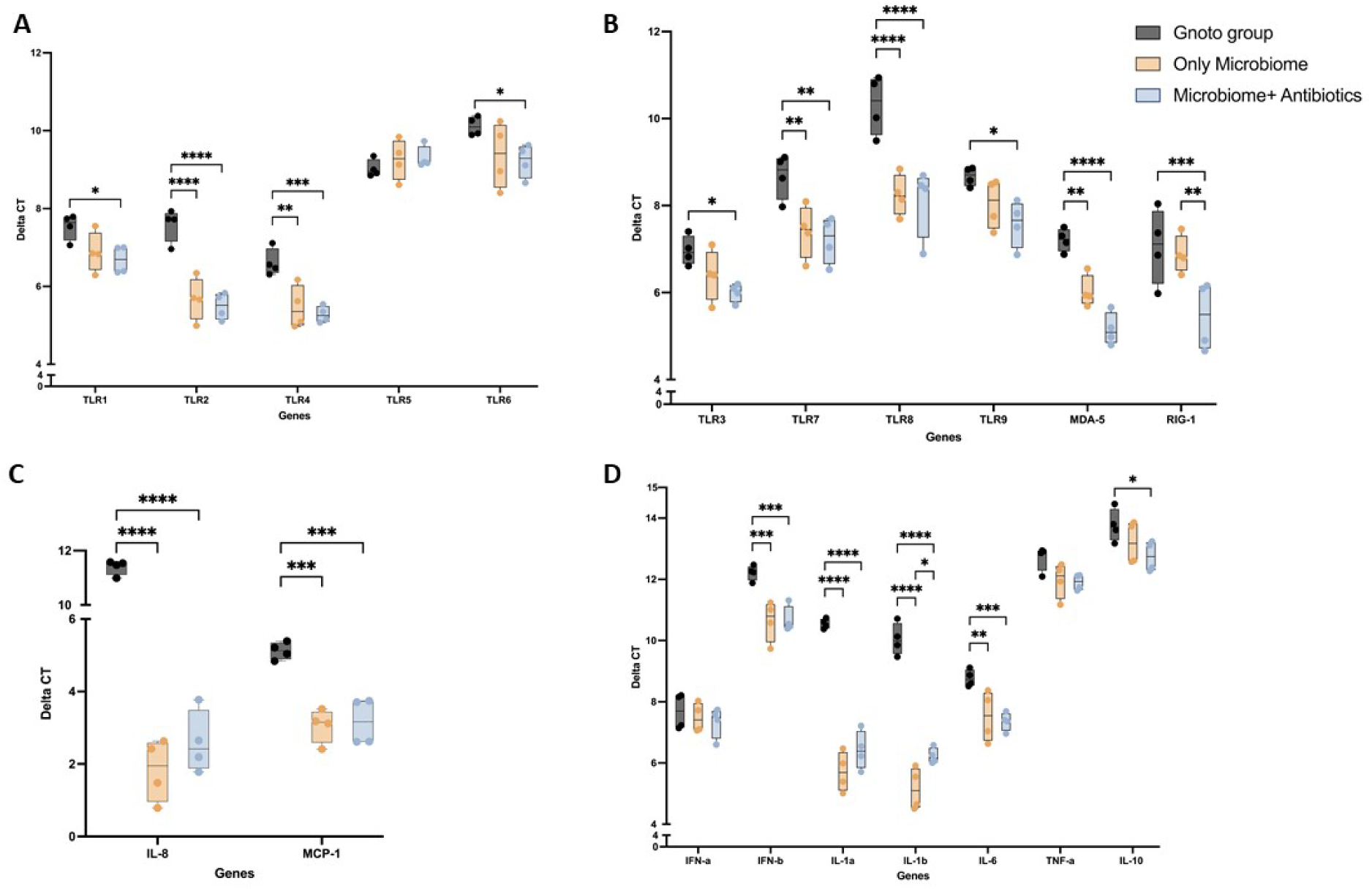
Gene expression profiles of cell surface TLRs (3a), intracellular TLRs and PRRs (3b), chemokines (3c), and cytokines (3d) in the nasal mucosa. ANOVA was applied to compare the means of delta cycle threshold (dCT), and upon observing significantly different expression levels, Tukey’s posthoc test was used. A P-value of less than 0.05 was considered statistically significant, and the error bar represents the standard error of the mean (SEM). The lower delta cycle threshold (dCT) value means a higher gene expression level. Ribosomal protein L4 gene (RPL4) was used as the reference gene to normalize the cycle threshold (CT) values.

TLR1 expression was notably higher in the antibiotics-treated group compared to controls. There were no significant differences in the expression of TLR5 and TLR6 between the groups.

Intracellular TLRs, particularly TLR3 and TLR9, were significantly upregulated in the antibiotics-treated group. Similarly, increased expression of TLR7 and TLR8 was observed in both nasal microbiome inoculated groups compared to the germ-free groups. Expression of MDA-5 was significantly higher in the antibiotics-treated group, followed by the microbiome-only inoculated group. RIG-I expression was also significantly higher in the antibiotics-treated group than in the other two groups. Regarding chemokines (Fig 3c), IL-8 and MCP-1 were significantly upregulated in both nasal microbiome inoculated groups.

For cytokines (Fig 3d), expressions of INF-β, IL-1α, and IL-6 were upregulated in both nasal microbiome inoculated groups. Expression of IL-1β was notably higher in the antibiotics group, followed by the microbiome-only inoculated group, compared to the germ-free control group. However, expressions of INF-α, TNF-α, and IL-10 did not show significant differences between the treatment groups.

Interestingly, in the mediastinal lymph nodes, expression of IL-1β and INF-β was downregulated in the nasal microbiome inoculated groups compared to the control germ-free group. TNF-α expression was also reduced in the antibiotics-treated group compared to the germ-free group.

The expression of other studied genes in mediastinal lymph nodes was found to be statistically non-significant between the groups. In the palatine tonsil, expression of TLR1 was significantly higher in the microbiome-only inoculated group than in the antibiotics-treated group. Expression of the remaining studied genes was non-significant between the groups. In the lungs, no significant changes in the expression level of the studied genes were detected between the treatment groups.

### Histopathology

Evaluation of H&E-stained nasal mucosa showed increase infiltration of lymphocytes, plasma cells and neutrophils in the microbiome inoculated groups. While microbiome inoculated group displayed blunted cilia and some mucosal erosion germ-free gnotobiotic group had thin mucosal barrier.

### Proliferation assay of PBMCs

PBMC were stimulated using ConA or LPS or left unstimulated. When stimulated with ConA, the proliferation of PBMC was higher in both nasal microbiome inoculated groups than in the control germ-free group (Fig 4). However, in LPS stimulated cells, proliferation was only higher in the microbiome plus antibiotics treated group and lower in the microbiome only inoculated group than in the control germ-free group. Though we detected changes in the proliferation activity, it was found to be statistically non-significant.

**Figure 4:**
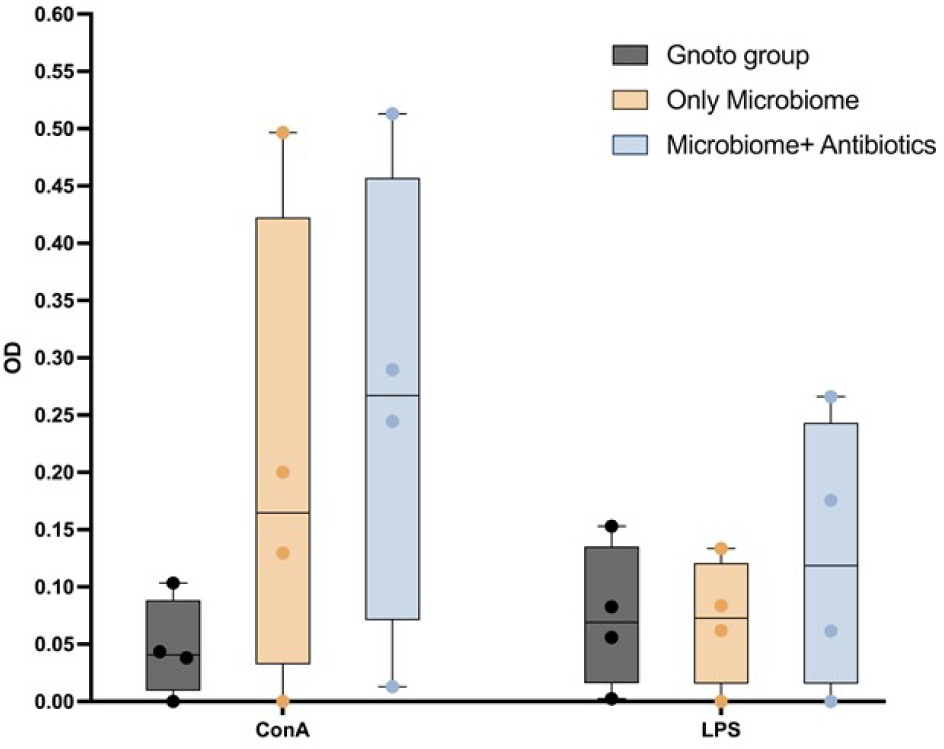
Proliferation activity of the PBMCs. Cells were stimulated with ConA or LPS or left untreated. BrdU, a substitute for nucleoside thymidine, was added. The incorporation of BrdU into proliferating cells was measured by substrate breakdown, and the readings were taken at 450nm wavelength. Data are expressed as the mean of three observations, error bar represents SEM. A p-value of less than 0.05 was considered statistically significant.

## Discussion

This study offers a detailed exploration of nasal microbiota transplantation in germ-free piglets, highlighting its impact on colonization across the respiratory and gastrointestinal tracts and subsequent effects on host immunity. Utilizing germ-free models enables a controlled analysis of these dynamics, devoid of confounding factors from pre-existing microbial communities.

One of the notable findings from this study is the successful colonization of the gastrointestinal tract by microbial taxa traditionally associated with the nasal cavity. Specifically, members of the phyla Proteobacteria and Firmicutes were observed to adapt to the gut environment, suggesting their potential as facultative anaerobes. Notably, families such as Clostridiaceae and Bacillaceae, which were significantly enriched in the gut post-inoculation, are also prevalent components of the healthy conventional pig gut microbiome[1, 2]. This presence indicates a functional integration of nasal-derived bacteria into the established gut ecosystem, emphasizing the adaptability of microbial communities.

The integration of these nasal-derived taxa into the gut highlights the potential interactions within the gut-lung axis, suggesting that microbial communities from one site may influence systemic health by modulating host immunity across both compartments. The presence of typical gut residents such as Clostridiaceae and Bacillaceae originating from the nasal microbiome underscores the complex interplay between different microbial communities and their roles in the host’s health.

The transplantation of nasal microbiota affected not only the microbial community structure but also host immune responses, as evidenced by the upregulation of various Toll-like receptors and changes in cytokine profiles. The involvement of gut-associated taxa such as Clostridiaceae and Bacillaceae, known for their immune-modulating properties in the gut, further supports the hypothesis that nasal microbiota can influence systemic immune functions.

Toll-like receptors are pattern recognition receptors (PRRs) and are an integral member of the innate immune system. These transmembrane proteins have leucine rich ectodomain which recognizes conserved pathogen associated molecular patterns (PAMPs) found on the pathogen. TLRs are expressed in a wide range of cells including innate and adaptive immune cells, epithelial cells, and non-immune cells such as fibroblasts. Activation of TLRs leads to production of pro-inflammatory cytokines, type I interferons and modulates the innate immune responses against invading pathogens (30, 31). In our study, in nasal mucosa we observed significant upregulation of TLR2, TLR4, TLR7, TLR8, and MDA-5 genes after nasal microbiome inoculation. Interestingly, expressions of TLR1, TLR3, TLR9, and RIG-1 were significantly upregulated only in antibiotics treated group. A similar pattern of expression of TLR1, TLR2, TLR3, TLR4, TLR5 and TLR6 was observed in nasal mucosa of conventional piglets at early age (32). In our study, we observed expression of MDA-5 was significantly higher in antibiotics treated group compared to both gnotobiotic and only nasal microbiome inoculated group. Interestingly, MDA-5 was also found to be significantly different between gnotobiotic and only nasal microbiome inoculated group. There was a decrease in the abundance of Moraxellaceae, Pasteurellaceae, Planococcaceae and Micrococcaceae families in nasal mucosa after the antibiotic treatment. Similarly, relative abundance of families Enterobacteriaceae, Weekellaceae, Lactobacillie and Corynebacteriaceae increased after the antibiotic treatment. This shift in the relative abundance of specific microflora following antibiotic administration may have induced alterations in the expression profiles of TLR1, TLR3, TLR9, and RIG-1 in only antibiotic treated group compared to both gnotobiotic and only nasal microbiome group. Furthermore, after the antibiotics treatment nasal microbiome composition shifted as visualized in PCA graph. The nasal microbiome diversity remained closely related to the inoculum in the non-antibiotic treated group. Similarly, LAD chart (Fig 1C) shows composition of highly abundance nasal microbiome after antibiotics treatment shifted from Pasteurellaceae, Streptococcaceae, Micrococcaceae, Alcaligenaceae to Mycoplasmataceae, and corynebactriaceae. Mycoplasma has an outer layer of lipoprotein layer which is detected by TLR1 (33) and interestingly, we observed increased abundance of mycoplasma after antibiotic treatment is concurrent with increased expression of TLR1 in the nasal mucosa.

This suggests a regulatory influence of nasal microbiome composition on the innate immune responses wherein depletion of enrichment of certain microbial communities by systemic antibiotics treatment triggers differential gene expressions of innate immune receptors (7, 34). These changes highlight the critical role in nasal microbiome in modulating host immune responses at mucosal surface potentially through PRRs, which are critical for recognizing pathogen molecular pattern and initiating early immune responses. This underscores the complex interplay between microbial diversity and host immune response at the respiratory mucosal surface (35–37). Overall, the composition of microbiome at the mucosal surface plays role in shaping early immune responses to invading pathogens and commensal microflora by altering expression of innate immune receptors (38).

Furthermore, we investigated if nasal microbiome colonization and antibiotics treatment influenced inflammatory pathways at the nasal mucosa. We studied expression of various cytokines and chemokines to observe innate immune responses generated by nasal microbiome colonization. We found increased expression of chemokines IL-8 and MCP-1 after nasal microbiome colonization. Similarly, we also observed increased expression of IL-1α, IL-1β, IL-6 and IFN-β following colonization of nasal microbiome. Simultaneously, we observed an infiltration of moderate number of lymphocytes, plasma cells and neutrophils in the underlying lamina propria of nasal mucosa by H&E staining (supplementary table1). This suggests nasal microbiome colonization mediated increase expression of chemokines and cytokines may have facilitate the recruitment of immune cells at the nasal mucosa. This underscores the importance of nasal mucosa in orchestrating complex inflammatory responses at the respiratory mucosal surface which could potentially determine the susceptibility and resistance of host to specific pathogens (32, 38).

We evaluated humoral immune responses to nasal microbiome colonization and quantified serum IgM, IgG and IgA levels. Previous study reported positive association between serum IgA levels and protection against rotavirus infection in gnotobiotic pig model (39). Similarly, experimental infection of PEDV in conventional pigs showed specific humoral IgA response after 14 days of inoculation. Although quantification of mucosal antibody levels was not performed in our study, reports from gnotobiotic pig study has substantiated serum IgA levels not IgG or IgM serve as reliable metric for protection against enteric and mucosal infections (39, 40). Consequently, significantly elevated serum IgA levels observed in nasal microbiome colonized groups indicates mucosal immune response generated by nasal microbiome colonization.

Subsequent to the colonization of respiratory mucosa by inoculated nasal microbiome unavoidably gut also got colonized. Notably, gut microbiome exhibited a predominance of Enterobacteriaceae and unlike nasal mucosa low abundance of Pasteutellaceae and Mycoplasmataceae. Intrestingly, fecal microbiome clustered more closely across all the animals (PCA graph) compared to nasal microbiome compared to nasal microbiome. Initially, at day 11 post inoculation gut microbiome alpha diversity was lower relative to nasal microbiome, suggesting that the gut microbiome may require a more extended period to achieve stabilization compared to relatively shorter nasal mucosa. Moreover, administration of the antibiotics appeared to induce a significant alteration in the composition of the gut microbiome as evidenced by increased abundance of Enterococcaceae and Lactobacillaceae. This suggests that antibiotic treatment may exert systemic effects and alter the microbiome composition on the mucosal surfaces across the body (41).

There are few limitations of our study, we did not evaluate the microbiome composition of lower respiratory tract and our swabbing technique only picks microbiome from upper respiratory tract. Depending on the diet, environment, stress, and health status upper respiratory tract microbiome composition can be transient and not stably colonized. Distinguishing the composition of upper and lower respiratory tract in gnotobiotic pig model can certainly aid our understanding on how microbes colonize and stabilize at different sites of respiratory tract and its effects in health and disease (42). Furthermore, the impacts of adoptively transferred sow serum on the colonization of the nasal microbiome were not comprehensively studied. Findings from our preliminary pilot study (unpublished) indicate that, in the absence of maternal antibody following nasal microbiome inoculation, piglets experienced severe illness. Mucosal surfaces were predominantly colonized by pathogenic microbes rather than commensals ones attributed to their undeveloped immune system. Absence of maternal antibody and subsequent colonization of pathogenic microbes from inoculum adversely influenced the piglets’ survival rates throughout the duration our preliminary study. Notably, there is a transfer of antibodies from sow to piglets via colostrum feeding in conventional swine farms, thereby suggesting that our method of administering sow serum to enhance piglet survival mirrors the principles of standard nursing care (43, 44).

In conclusion, we established a gnotobiotic pig model to study the dynamics of nasal microbiome colonization and its consequential influence on the activation of innate immune responses. Nasal microbiome has been traditionally overshadowed by gut microbiome in the context of studying its contributions in mucosal infection. Our study provides a comprehensive analysis of the nasal microbiome, the subsequent innate immune responses and the impact of antibiotic administration on both the microbiome and immune function. Further research is imperative to elucidate the complex molecular mechanisms through which the nasal microbiome influences the adaptive and innate immune system thereby affecting susceptibility to mucosal infections.

---

**Supplementary figure 1.**
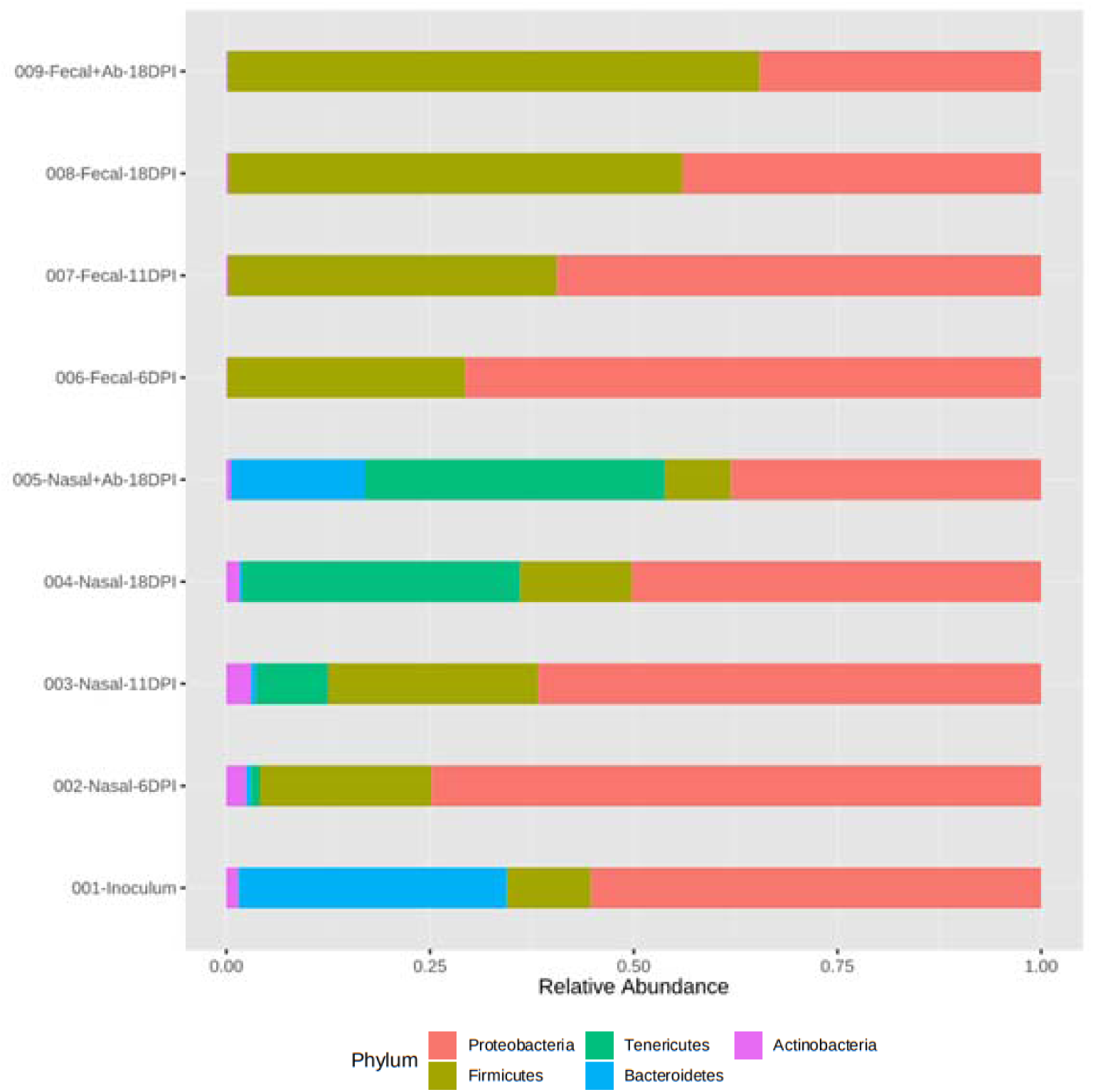
Relative abundance of bacterial taxa at phylum level. Stacked bar plot shows the relative abundance of major top 5 phyla in the nasal and fecal community collected at different time points. Inoculum represents the pooled donor nasal microbiota community. Color legend represents the phylum.

**Supplementary figure 2.**
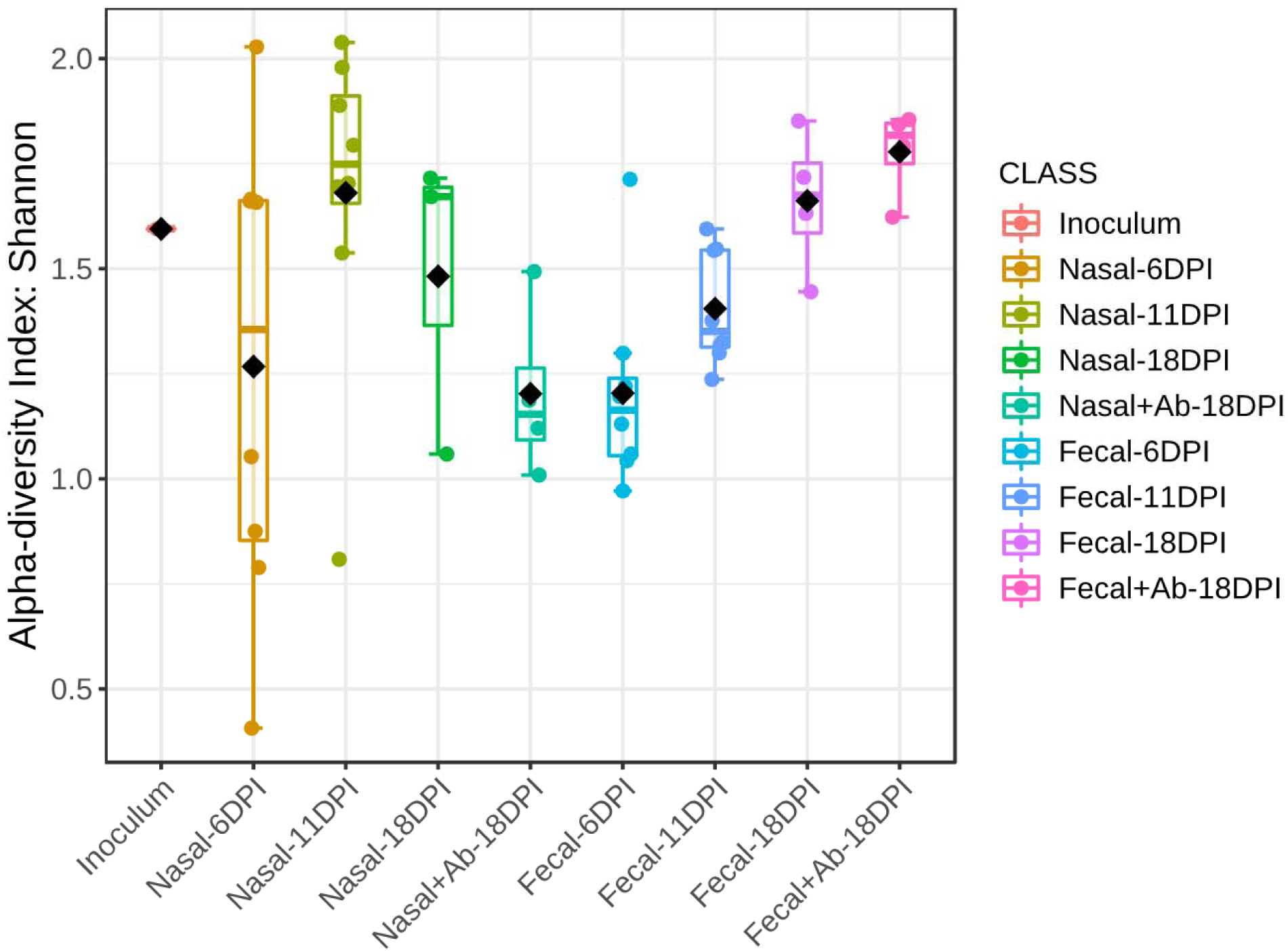
Alpha diversity measurement of nasal and fecal community after nasal microbiota transplantation. Box plots represents the Shannon alpha diversity index measured for individual samples at different time points. Inoculum represents the pooled conventional piglets’ nasal microbiota.

**Supplementary figure 3.**
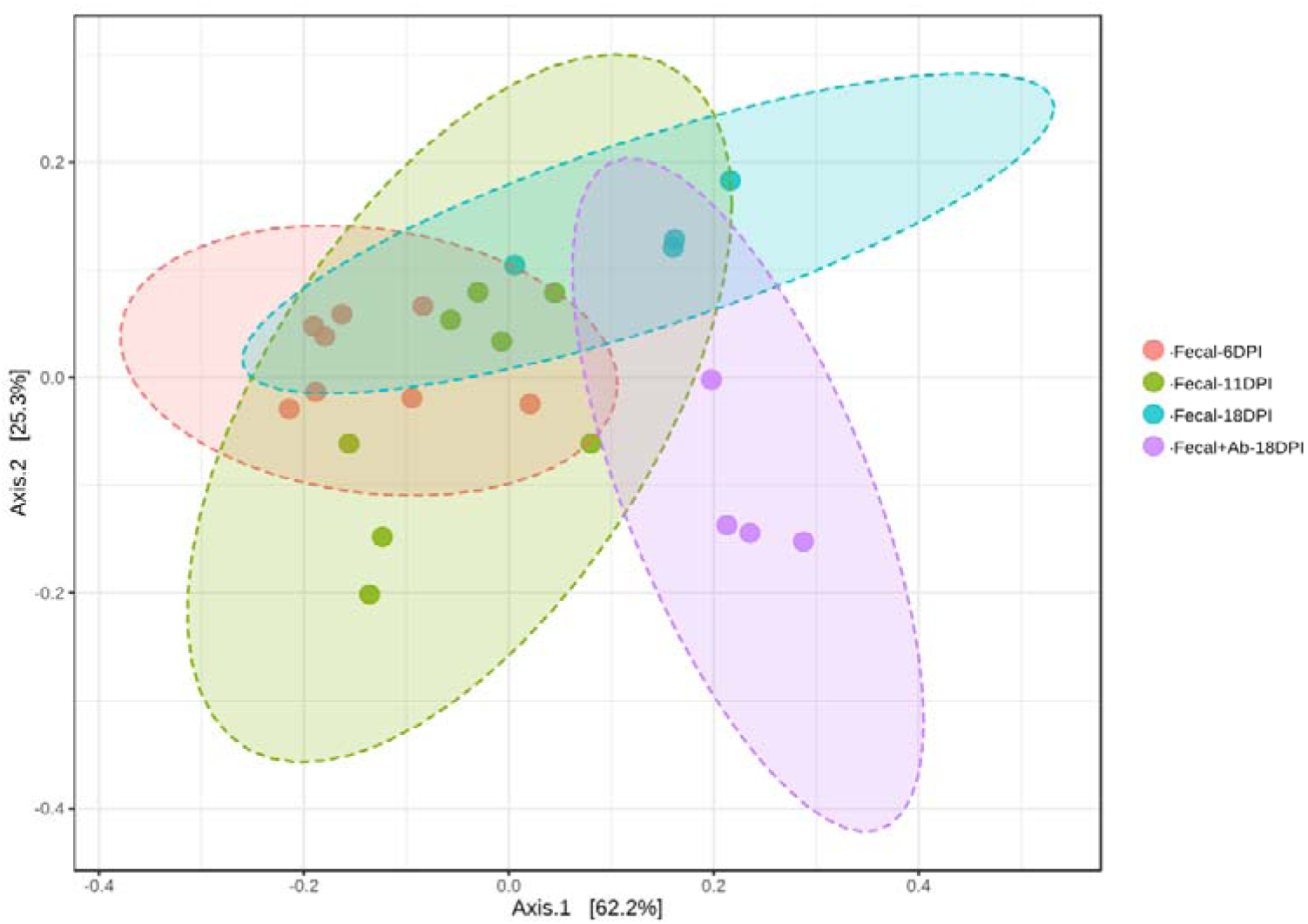
Beta diversity measurement of fecal community of gnotobiotic piglets after nasal microbiota transplantation. PCoA plot shows the clustering of fecal samples based on their Bray Curtis dissimilarity index at significance of p-value <0.001 (PERMANOVA).

